# Leptospiral lipopolysaccharide dampens inflammation through upregulation of autophagy adaptor p62 and NRF2 signaling in macrophages

**DOI:** 10.1101/2022.08.24.505079

**Authors:** Delphine Bonhomme, Ignacio Santecchia, Pedro Escoll, Stylianos Papadopoulos, Frédérique Vernel-Pauillac, Ivo G. Boneca, Catherine Werts

## Abstract

*Leptospira interrogans* are pathogenic bacteria responsible for leptospirosis, a worldwide zoonosis. All vertebrates can be infected, and some species like humans are susceptible to the disease whereas rodents such as mice are resistant and become asymptomatic renal carriers. Leptospires are stealth bacteria that are known to escape several immune recognition pathways and resist killing mechanisms. We recently published that leptospires may survive intracellularly and exit macrophages, in part by escaping xenophagy, a pathogen-targeting form of autophagy. Interestingly, autophagy is one of the antimicrobial mechanisms often highjacked by bacteria to evade the host immune response. In this study we therefore explored whether leptospires subvert the key molecular players of autophagy to facilitate the infection. We showed in macrophages that leptospires triggered a specific accumulation of autophagy-adaptor p62 in puncta-like structures, without major alteration of autophagy flux. Unlike active bacterial mechanisms described to date, we demonstrated that leptospires trigger p62 accumulation using a passive mechanism of LPS signaling via TLR4/TLR2. p62 is a central pleiotropic protein, not only involved in autophagy, but also mediating cell stress and death, *via* the translocation of transcription factors. We demonstrated that *Leptospira*-driven accumulation of p62 induced the translocation of transcription factor NRF2. However, NRF2 translocation upon *Leptospira* infection did not result as expected in antioxydant response, but dampened the production of inflammatory mediators such as iNOS/NO, TNF and IL6. Overall, these findings highlight a novel passive bacterial mechanism linked to p62/NRF2 signaling that decreases inflammation and contributes to the stealthiness of leptospires.

## INTRODUCTION

*Leptospira interrogans* are spirochete bacteria and the causative agent of leptospirosis, a neglected worldwide zoonosis whose global impact on human health is increased due to climate change, and causes around 60 000 deaths *per* year worldwide (1). Although all vertebrates can be infected by leptospires, they do not all present the same symptoms and susceptibility to the disease (2). Humans, as sensitive hosts, may suffer from acute leptospirosis ranging from flu-like symptoms to multi-organ failure in 5-10% cases (1). On the other hand, rodents such as mice and rats are resistant to acute illness and become chronically colonized upon infection (3). Leptospires establish a stable colonization in the proximal tubules in the kidneys, that leads to their excretion in the urine throughout the life of the animal, contributing to the spread of the zoonosis (3, 4).

Upon infection, the host immune defenses rely on humoral and cellular components, such as the complement system, immunoglobulins, and patrolling phagocytes. Activation of immune cells such as macrophages is mediated by the sensing of microbial associated molecular patterns (MAMPs) by pattern recognition receptors (PRRs). Leptospires are agonists of TLR2, through their numerous lipoproteins (5, 6), and activate the NLRP3 inflammasome (7). However, they are remarkable as stealth pathogens that escape recognition by NOD receptors (8) and by TLR5 (9) through unique mechanisms. Furthermore, the leptospiral lipopolysaccharide (LPS), a central virulence factor (10), possesses structural peculiarities (11) that do not activate human TLR4 (12), whereas they partially activate murine TLR4 (13). In addition to escaping PRR recognition, leptospires also escape some phagocytic functions. Although leptospires are mostly extracellular bacteria, we previously reported that they can be found within macrophages without intracellular replication (14), and are neither targeted by phagocytosis nor classical microbicidal compounds (14). We further excluded that intracellular leptospires could be targeted by xenophagy (14), a specific form of autophagy promoting the degradation of intracellular pathogens (15–17).

Interestingly, among the numerous proteins involved in autophagy, many have pleiotropic roles that are not restricted to autophagic-degradation and are at the crossroads between autophagy, cell death, cellular stress and inflammation. For instance, autophagy adaptors p62 and NDP52, which traditionally bridge the cargo and autophagosome for specific degradation, have been associated with many xenophagy-independent inflammatory modulations (18, 19). Both p62 and NDP52 have been shown to mediate nuclear translocation of transcription factors NFκB and NRF2, which are involved in inflammation and cellular stress, respectively (18, 20).

In this study, we focused our investigations on autophagy adaptors and unexpectedly found in macrophages infected with leptospires a role of their LPS in dampening of inflammation through a modulation of the p62/ NRF2 axis.

## MATERIALS AND METHODS

### Leptospira interrogans cultures

*Leptospira interrogans* used (Manilae strain L495, Copenhageni strain Fiocruz L1-130, Icterohaemorragiae strain Verdun) and *Leptospira biflexa* (Patoc strain Patoc I) were grown in Ellinghausen-McCullough-Johnson-Harris (EMJH) medium at 28°C without agitation and diluted weekly (twice a week for *L. biflexa*) to obtain exponential cultures. For infection, cultures were centrifuged (3250 g, 25 min), resuspended in PBS (Lonza), and enumerated in Petroff-Hauser chamber. Inactivated leptospires (“heat-killed”) were heated at 56 °C for 30 min with mild agitation. For fluorescent labelling, 10 mL of exponential culture was centrifuged (3250 g, 25 min) and the pellet was resuspended in the same volume of PBS (Lonza) with addition of 10 µM CFSE (Sigma) for 30 min, followed by one wash in PBS before counting and infection.

### Mice experiments

Adults male and female C57/BL6 mice (Janvier Labs, Le Genest-Saint-Isle, France) were injected *via* intraperitoneal route with 1 x10^8^ heat-killed leptospires/mouse in PBS. Next, 4h post-infection mice were euthanized, and peritoneal content was recovered as previously described (21). Peritoneal cells were plated at 1 x10^6^ cells/mL and were let to adhere for 1h before fixation and immunofluorescence staining.

### Cell culture

BMDMs were obtained and derived from adults male and female mice of either C57BL/6 WT (Janvier Labs, Le Genest-Saint-Isle, France) or TLR2^-/-^ TLR4^-/-^ (from Institut Pasteur) as described before (13). BMDMs were seeded in plates (TPP) the day before infection at a concentration of 0.8 x10^6^ cells/mL, in antibiotic-free complete RPMI (RPMIc) (containing glutamine (Lonza), supplemented with 10 % v/v heat-inactivated fetal calf serum (HI-FCS, Gibco), 1 mM non-essential amino acids (NEA, Gibco) and 1 mM sodium pyruvate (NaPy, Gibco)).

RAW264.7 murine macrophage-like cell line was cultured in antibiotic-free RPMIc. RAW-mLC3-DiFluo (Invivogen) autophagy reporter cells (transfected with LC3-GFP-RFD) were cultured with 100 µg/mL zeocin (Invivogen). Cells were seeded in plates (TPP) the day before infection at a concentration of 0.3 x10^6^ cells/mL, in antibiotic-free RPMIc.

Human THP-1 monocyte-like cell line, stably transfected with CD14 (THP1-CD14), was cultured in RPMIc and seeded at 1 x10^6^ cells/mL.

All cell cultures were tested negative for *Mycoplasma* contamination, and all cell cultures used for experiments were maintained under 80 % confluence in order to prevent autophagic stress.

Cell starvation was induced in EBSS media (Gibco) after 2 washes. Autophagy blockage was induced with 100 nM bafilomycin A1 (BafA1, Sigma-Aldrich) 4h before cell collection. Inflammasome inhibition was triggered using 25 µM glibenclamide (Thermo Fisher) 30 min before infection. Finally, stimulations by leptospiral LPS were performed with 1 µg/mL of LPS of *L. interrogans* L495 extracted from the phenolic phase of the hot water/phenol extraction protocol, as we recently reviewed (22).

### Immunofluorescence and High Content (HC) automated microscopy

Cells were seeded and infected on cover glass (18 mm diameter, # 1.5 thickness, Electron Microscopy Science) and were fixed in 4 % v/v *para*-formaldehyde for 10 min, followed by three washes with PBS. Blocking was 1h in PBS + 5% w/v bovine serum albumin (BSA) and 2.5 µg/mL anti-CD16-CD32 (FcBlock, Thermo Fisher). Primary antibodies (**Table 1**) or isotypes were incubated overnight at 4 °C in PBS + 1% w/v BSA, 0.05% w/v saponin. Cells were then washed three times with PBS + 0.05% w/v saponin and labeled for 1h with secondary antibodies when necessary (**Table 1**) and 1µg/mL DAPI in PBS + 1% w/v BSA, 0.05% w/v saponin. For NRF2 nuclear staining, permeabilization was performed in PBS + 1% w/v BSA, 0.5% v/v Triton X-100 for 10 min prior to blocking. Image acquisition was performed on Leica SP5 confocal microscope (63x - 1.4 NA, oil immersion). Default settings were used and both UV and argon lasers were used at power between 10-30 %.

**Table 1.** Antibodies list for immunofluorescence and Western blot analyses

| Application* | Target | Clonality** | Conjugated | Host | Dilution | Reference | Supplier |
| --- | --- | --- | --- | --- | --- | --- | --- |
| IF | **p62** | mAb | No | Rabbit | 1:500 | ab109012 | Abcam |
|  | Rabbit IgG | pAb | AlexaFluor 488 | Goat | 1:1.000 | A-11034 | ThermoFisher |
| IF | **Nrf2** | mAb | No | Rat | 1:250 | #14596 | CST |
|  | Rat IgG | pAb | AlexaFluor 647 | Goat | 1:1.000 | A-21247 | ThermoFisher |
| IF | **F4/80** | mAb | APC-Cyanine7 | Rat | 1:250 | 123117 | BioLegend |
| WB | **LC3** | pAb | Purified | Rabbit | 1:1.000 | L7543 | Sigma |
|  | **p62** | mAb | Purified | Mouse | 1:1.000 | ab109012 | Abcam |
|  | **OPTN** | pAb | Purified | Rabbit | 1:1.000 | ab23666 | Abcam |
|  | **NDP52** | pAb | Purified | Rabbit | 1:1.000 | GTX115378 | GeneTex |
|  | Rabbit IgG | pAb | HRP | Goat | 1:10.000 | 7074S | CST |

\* IF: immunofluorescence, WB: Western blot, \*\* mAb: monoclonal antibody, pAb: polyclonal antibody

For HC automated microscopy, cells were seeded in dark, transparent bottom 96-well plates (Greiner, µClear) at 0.1 x10^6^ cells/mL. Cells were stained as described, except for RAW-mLC3-DiFluo that were fixed 10 min in cold ethanol/acetone (1:1). Imaging was performed on Opera Phenix (Perkin Elmer) using confocal settings and 63X water-immersion objective (NA 1.15). Light source was kept at 20 to 50% power and exposure time was set to obtain intensities 1000-5000. Automated acquisition allowed analysis of 500-1000 cells/well in technical triplicates for each experiment. Automated image analysis workflow and puncta quantification was performed using Columbus image data storage and analysis system (PerkinElmer), with the following steps: *(i)* import data; *(ii)* find nuclei with DAPI signal; *(iii)* find cytoplasm using target protein signal (p62 / NRF2 / F4/80); *(iv)* select cell population of interest if relevant (F4/80^+^ peritoneal cells), *(v)* find spots if relevant (p62); *(vi)* formulate classical outputs: number of p62 puncta, NRF2 intensity in nucleus/cytoplasm; *(vii)* formulate calculated outputs: %p62 positive cells, NRF2 ratio, %NRF2 positive cells*; (viii)* save script and run batch analysis and *(ix)* export data.

### SDS-PAGE and Western blot

For Western blot, cells were scrapped and centrifuged (400 g, 10 min). Pellets were resuspended in 50 µL of ice-cold RIPA lysis buffer supplemented with 1x complete Mini, EDTA-free proteases inhibition cocktail (Roche). Cells were lysed on ice for 15 min followed by centrifugation (12000 g, 40 min, 4 °C). Soluble proteins were recovered in the supernatant and dosed using the Bradford assay and the samples were denatured in Laemli buffer (99 °C, 10 min). SDS-PAGE was performed at 100 V on 4-15% gradient acrylamide Stain-free gels in Tris-Glycine-SDS buffer (BioRad), with 5-10 µg of protein. Total proteins were visualized as internal control using stain free reagent in ChemiDoc (BioRad) with a 5 min exposition time and then transferred on 0.22 µm PVDF membrane (BioRad) using Mixed MW fast transfer of 1 miniGel (BioRad). Membrane was blocked 1h in TBS with 0,05% v/v Tween 20 (TBS-T) + 5% w/v BSA. The membrane was probed overnight at 4 °C with primary antibodies (**Table 1**) in TBS-T + 5% w/v BSA. After three washes, membrane was incubated 1h with secondary anti-rabbit IgG HRP-linked (**Table 1**) in TBS-T + 5% w/v BSA. After three washes, blots were revealed using the Clarity reagent (BioRad) with automatic exposure time.

### Small interfering RNA transfection

For siRNA transfection, cells were seeded at confluence of 0.1 x10^6^ cells/mL the day before transfection. Transfection was performed using Lipofectamine siRNA Max reagent (ThermoFisher) OptiMEM media (Gibco) and according to the manufacturer’s instructions. The final concentration of commercial predesigned siRNA targeting *p62* and *nrf2* (FlexiTube, Qiagen) was 80 nM and the corresponding scramble siRNA (all start negative, Qiagen) was used as control. The siRNA preparations in lipofectamine were incubated at room temperature for 25 min before delicate transfection of the cells in OptiMEM. After 8 hours, the same volume of RPMIc was added for overnight incubation. Infection was performed the next day in RPMIc after complete media removal.

### ELISA and Griess reaction

Fresh cell culture supernatants were used 24h post-infection for dosage of nitric oxide (NO) by the Griess reaction. Cytokine dosage was performed on frozen supernatants by ELISA using DuoSet kits (R&D Systems), according to the manufacturer’s instructions. NO and cytokine concentrations were plotted both before and after normalization by cell viability.

### Viability and LDH release assays

LDH release was measured using the CyQuant LDH Cytotoxicity Assay kit (fresh supernatant) (Invitrogen), according to the manufacturers’ instructions. Viability was measured using MTT assay: cells were incubated for 2h in 1 mg/mL MTT solution (Sigma) in complete RPMI. MTT crystals were then dissolved in HCl 1 M, isopropanol (1:24) and absorbance was read at 595 nm. For each technical replicate, the viability value was used to normalize both NO and cytokines measurements using the following formula: 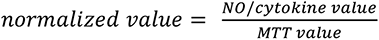.

### RT-qPCR

RNAs were extracted from frozen cells using RNeasy Mini Kit (Qiagen) according to the manufacturer’s instructions. cDNAs were then obtained by retro-transcription (RT) using SuperScript II RT (Invitrogen). The equivalent of 20ng of cDNA were used for RT-qPCR with specific primers and probes (**Table 2**). qPCR was performed using Taqman Universal MasterMix (Applied Biosystems) on a StepOne Plus Real Time PCR System (Applied Biosystems) with the standard protocol. Fold changes were calculated with the 2^-ΔΔC^_T_ method, using hypoxanthine guanine phosphoribosyl transferase (HPRT) as internal control.

**Table 2.**
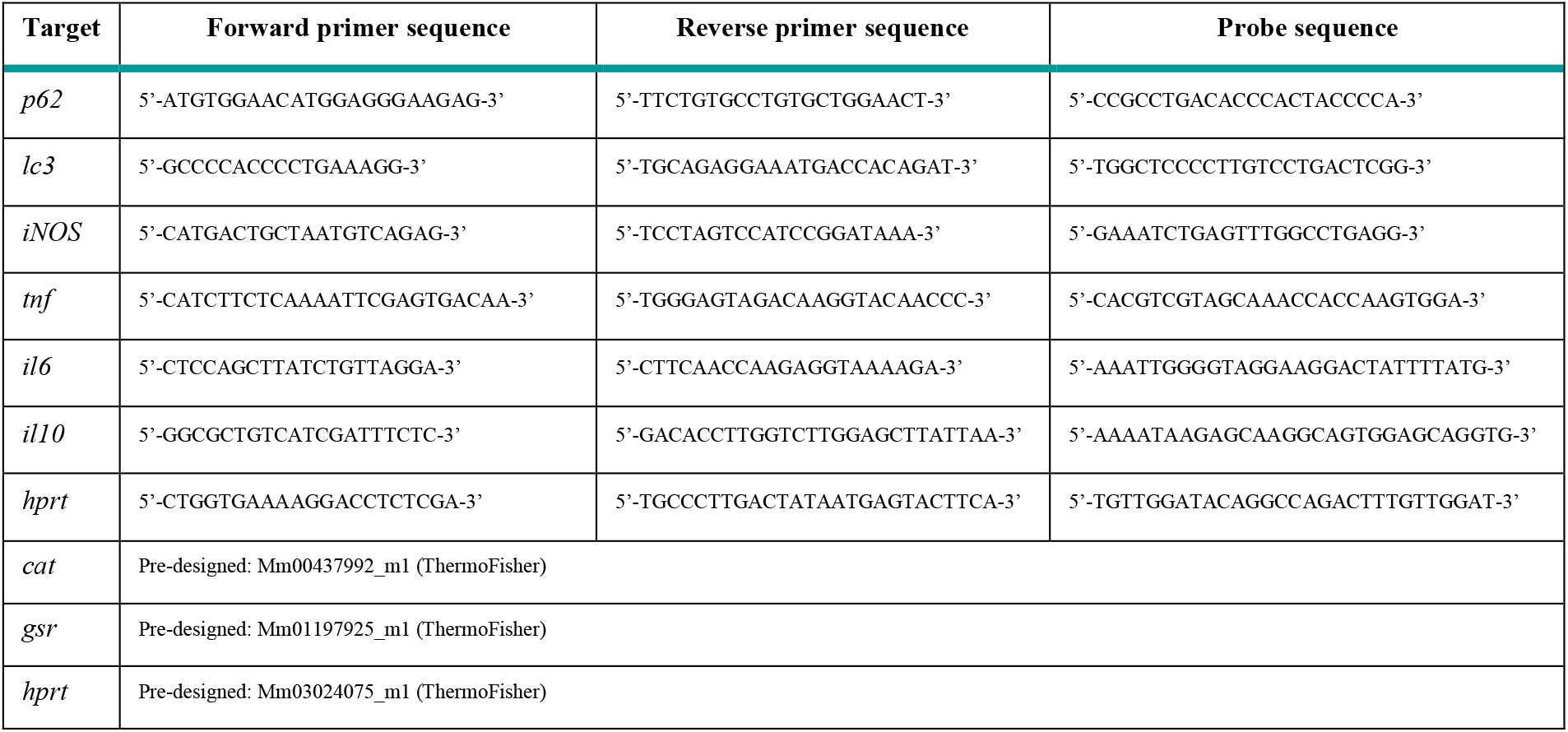
Primers and probe list for RT-qPCR analyses

### Statistical analyses

All statistical analyses were performed using Student’s *t*-test with corresponding *p* values: * for *p* < 0.05; ** for *p* < 0.01 and *** for *p* < 0.001.

## RESULTS

### *L. interrogans* induces upregulation and specific accumulation of autophagy-adaptor p62 in puncta-like structures

To address modulation of autophagy-adaptors upon infection with leptospires, we infected bone marrow derived macrophages (BMDMs) with *L. interrogans* Manilae strain L495 and analyzed by Western blot the levels of the different adaptors p62, NDP52 and Optineurin. We observed an accumulation of p62 over time upon infection (**Fig 1A**). We confirmed this phenotype using other pathogenic serotypes of *L. interrogans*: Copenhageni strain Fiocruz L1-130 and Icterohaemorrhagiae strain Verdun, and using the saprophytic *Leptospira biflexa* Patoc strain Patoc I (**Fig 1B**). Epifluorescence analyses of BMDMs infected 4h with *L. interrogans* further revealed that infection triggers accumulation of p62 in puncta-like structures (**Fig 1C**). This phenotype seemed specific of p62 since we did not observe accumulation of the autophagy adaptors (NDP52 & optineurin) upon leptospira infection (**Sup. Fig 2A**).

**Figure 1.**
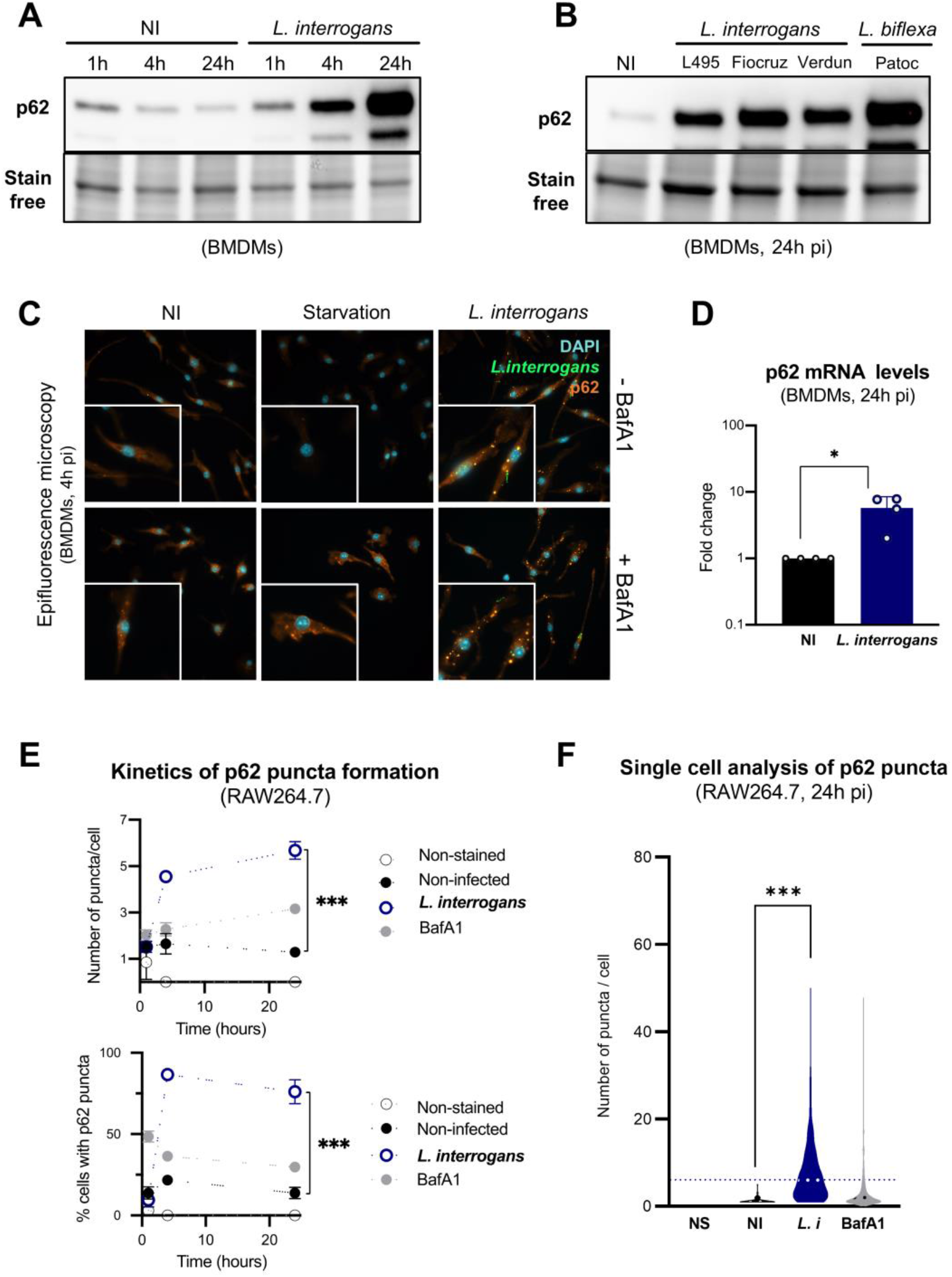
*L. interrogans* induces upregulation and specific accumulation of autophagy-adaptor p62 in puncta-like structures. **A)** WB of p62 in BMDMs 1h, 4h and 24h post-infection with *L. interrogans* serovar Manilae strain L495 at MOI 100. Gel stain free is presented as loading control of total proteins. **B)** WB of p62 in BMDMs 24h post-infection with *L. interrogans* serovar Manilae strain L495, serovar Copenhageni strain Fiocruz L1-130, serovar Icterohaemorrhagiae strain Verdun, and *L. biflexa* serovar Patoc strain Patoc I, at MOI 100. **C)** Epifluorescence analyses of p62 in BMDMs 4h post-infection with fluorescently-labeled *L. interrogans* serovar Manilae strain L495 at MOI 10, or 4h upon starvation induction in EBSS medium, with or without 100nM of bafilomycin A1 (BafA1). **D)** RT-qPCR analyses of p62 mRNA levels in BMDMs 24h post-infection with *L. interrogans* serovar Manilae strain L495 at MOI 100. Bars correspond to mean +/- SD of technical replicates (*n*=4). **E)** Automated confocal microscopy analyses of the number of p62 puncta *per* cell and the percentage of p62 positive cells in RAW264.7 cells 1h, 4h and 24h post-infection with *L. interrogans* serovar Manilae strain L495 at MOI 100, or stimulation with 100nM of BafA1. Dots correspond to mean +/- SD of technical replicates (*n*=3). **F**) Single cell microscopy analysis of the number of p62 puncta *per* cell 24h post-infection of RAW264.7 cells with *L. interrogans* serovar Manilae strain L495 at MOI 100, or stimulation with 100nM of BafA1. Dashed lines correspond to median. **A-F)** Data presented are representative of at least 3 independent experiments. Statistical analyses were performed using Student’s *t*-test with corresponding *p* values:* for *p* < 0.05 and *** for *p* < 0.001.

Among other mechanisms, autophagy adaptors are constitutively degraded by autophagy upon autophagosome / lysosome fusion. We hypothesized that p62 accumulation could be triggered by blockage of autophagy flux by leptospires. However, when we monitored the autophagy hallmark protein LC3-II in BMDMs by Western blot, no LC3-II accumulation was observed upon *Leptospira* infection, in contrast to Bafilomycin (BafA1) treatment that blocks the autophagy flux, causing LC3-II to accumulate (**Sup. Fig 1A**). Thus we conclude that infection with leptospires does not alter the autophagy flux. Unexpectedly, we observed a mild reduction in LC3-II levels upon infection (**Sup. Fig 1B**). Using murine macrophages RAW-mLC3-diFluo cells analyzed by automated microscopy, we were able to confirm that leptospires induce a mild decrease of the number of autophagosomes without altering the number of autolysosomes, again showing no alteration of the autophagy flux (**Sup. Fig 1C-D**).

We then analyzed the transcriptional regulation of p62 in BMDMs by RT-qPCR and observed a significant upregulation of p62 mRNA 24 h post-infection with *L. interrogans* (**Fig 1D**). This corroborated the idea that p62 accumulation is not mediated by autophagy blockage. The kinetics of the p62 puncta formation were further characterized in RAW264.7 cells, using automated high content (HC) confocal microscopy. Results showed time-dependent increase of both the number of p62 puncta *per* cell and the percentage of p62 positive cells (**Fig 1E** **& Sup. Fig 2B**). Single cell analysis confirmed an average of 5-10 puncta *per* cell and highlighted cell-to-cell heterogeneity, with some macrophages containing up to 60 puncta (**Fig 1F**). Of note, BafA1 that blocks autophagy flux also led to p62 accumulation as expected, but to a much lower extent than infection with leptospires (**Fig 1C, 1E & 1F**). Altogether, these results confirm our previous results showing that *L. interrogans* do not induce autophagy in murine macrophages (14), and suggest that leptospires induce a specific accumulation of p62, not linked to an autophagy blockage.

### p62 accumulation is triggered by the leptospiral LPS through TLR4 & TLR2

Many bacteria interfere actively with autophagy molecules *via* secreted effectors or RNA interference (16, 23–28). To investigate whether such active mechanisms are also used by leptospires, we analyzed BMDMs and RAW264.7 cells after stimulation with heat-killed (HK) leptospires. We observed p62 accumulation visible by Western blot (**Fig 2A**) and quantified by automated microscopy (**Fig 2B** **& 2C**). These findings exclude an active leptospiral mechanism and suggest a role for the recognition of leptospiral MAMPs. We therefore stimulated cells for 24h with leptospiral LPS and observed that such stimulation recapitulates p62 accumulation (**Fig 2A, 2B & 2C**). As p62 accumulation seemed to be mediated by recognition of the leptospiral LPS, we investigated the roles of TLR4 & TLR2. In murine cells, these two receptors are respectively activated by the leptospiral lipid A, and the leptospiral lipoproteins that co-purify with the LPS (5, 12). We infected WT and TLR2/4 dko BMDMs with *L. interrogans* for 24h, and analyzed by Western blot and automated microsopy. We observed that p62 accumulation was greatly dampened in TLR2/TLR4 dko cells (**Fig 2D** **& 2E**). Finally, we analyzed mRNA levels by RT-qPCR and observed that the increase in p62 mRNA levels observed in WT BMDMs was abolished in TLR2/4 dko BMDMs (**Fig 2F**). Overall, these data show that p62 accumulation is mediated by TLR2/4 recognition of the leptospiral LPS. To address if such mechanism was conserved *in vivo*, we injected C57BL/6 mice intraperitoneally with 1 x10^8^ heat-killed *L. interrogans*, representative of MAMPs and potent TLR2/TLR4 agonists, that present the advantage of not replicating nor disseminating upon injection. Interestingly, using automated microscopy we showed that p62 was also accumulated *in vivo* in peritoneal F4/80^+^ macrophages (**Fig 2G**).

**Figure 2.**
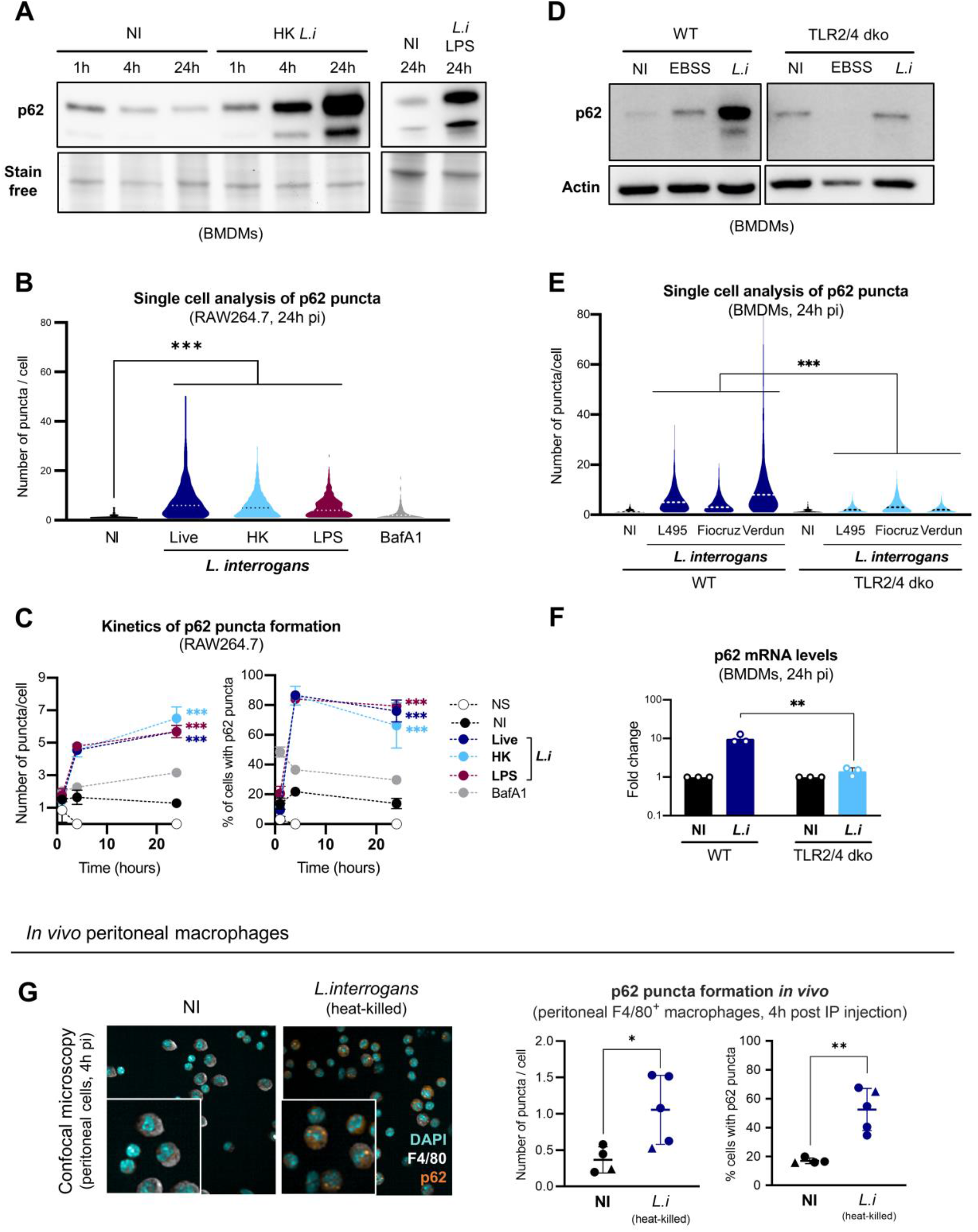
p62 accumulation is triggered by the leptospiral LPS through TLR4 & TLR2. **A)** WB of p62 in BMDMs 1h, 4h and 24h post-stimulation with heat-killed (56°C, 30 min) *L. interrogans* serovar Manilae strain L495 at MOI 100 or 24 h post-stimulation with 1 µg/mL of the corresponding purified leptospiral LPS. Gel stain free is presented as a loading control of total proteins. **F)** Single cell microscopy analysis of the number of p62 puncta *per* cell 24h post-infection of RAW264.7 cells with either live or heat-killed (56°C, 30 min) *L. interrogans* serovar Manilae strain L495 at MOI 100, stimulation with 1 µg/mL of the corresponding purified leptospiral LPS, or stimulation with 100nM of bafilomycin A1 (BafA1). Dashed lines correspond to median. **C)** Automated confocal microscopy analyses of the number of p62 puncta *per* cell and the percentage of p62 positive cells 1h, 4h and 24h post-stimulation of RAW264.7 cells with MOI 100 of either live or heat-killed (56°C, 30 min) *L. interrogans* serovar Manilae strain L495, stimulation with 1 µg/mL of the corresponding purified leptospiral LPS, or stimulation with 100nM of BafA1. Dots correspond to mean +/- SD of technical replicates (*n*=3). **D)** WB of p62 in WT and TLR2/4 dko BMDMs 24h post-infection with *L. interrogans* serovar Manilae strain L495 at MOI 100, or upon starvation induction in EBSS medium. WB of actin is presented as loading control. **E)** Single cell microscopy analysis of the number of p62 puncta *per* cell 24h post-infection of WT and TLR2/4 dko BMDMs with *L. interrogans* serovar Manilae strain L495, serovar Copenhageni strain Fiocruz L1-130 or serovar Icterohaemorrhagiae strain Verdun at MOI 100. Dashed lines correspond to median. **F)** RT-qPCR analyses of p62 mRNA levels in WT and TLR2/4 dko BMDMs 24h post-infection with *L. interrogans* serovar Manilae strain L495 at MOI 100. Bars correspond to mean of independent experiments (*n*=3). **G)** Automated confocal microscopy images and analyses of the number of p62 puncta *per* cell and the percentage of p62 positive cells in adherent F4/80^+^ peritoneal cells 4h post intra-peritoneal injection of 1 x10^8^ heat-killed (56°C, 30 min) *L. interrogans* serovar Manilae strain L495 in C57BL/6J mice. Dots correspond to individual mice from 3 independent experiments (round symbols = females / triangle symbols = males). **A-C, F, G)** Data presented are representative of at least 3 independent experiments. **D-E)** Data presented are representative of 2 independent experiments. Statistical analyses were performed using Student’s *t*-test with corresponding *p* values: * for *p* < 0.05; ** for *p* < 0.01 and *** for *p* < 0.001.

Finally, we ask whether this mechanism of p62 accumulation could be conserved in human cells. Indeed, there is a TLR4 host species-specificity of the innate immune recognition of leptospires (29). Leptospiral lipid A activates mouse-TLR4 but not human-TLR4 (12). However, lipoproteins that co-purify with the LPS activate both mouse-TLR2 and human-TLR2 in a CD14-dependent manner (12). We therefore infected human monocytic THP1-CD14 cells with *L. interrogans* serovar Manilae strain L495. We also observed a specific p62 accumulation and no NDP52 accumulation after infection with leptospires (**Sup. Fig 2C**), suggesting conserved mechanism of p62 accumulation in human and murine macrophages, and a prominent role of TLR2 in human cells.

Leptospires have been shown to activate the NLRP3 inflammasome, in humans (30, 31), and in a TLR2/4-dependent manner in mice (7). Furthermore, NLRP inflammasomes have been shown to modulate autophagy (32–35). Therefore, we asked whether activation of inflammasome could play a role in p62 accumulation and LC3-II diminution. We stimulated BMDMs with both live and heat-killed leptospires in the presence or absence of NLRP3 inhibitor glibenclamide. Efficiency of the treatment was controlled by measuring IL1β production after 24h (**Sup. Fig 3A**). p62 and LC3-II levels were analyzed by Western blot. We observed a similar levels of p62 accumulation upon infection with either live or heat-killed leptospires in both control and glibenclamide treated cells (**Sup. Fig 3B**). Consistently, the diminution of LC3-II upon infection was visible in both control and glibenclamide treated cells (**Sup. Fig 3B)**. Overall, this suggests that the modulation of autophagy players by leptospires, although TLR2/4-dependent, is not mediated by activation of the NLRP3 inflammasome.

### Leptospires and their LPS trigger translocation of transcription factor NRF2

Stress pathways are induced when autophagy adaptors accumulate in the cell (i.e. in the absence of functional autophagy or because of specific upregulation). p62 accumulation induces translocation of stress-responsive nuclear factor erythroid 2-related factor 2 (NRF2) (20, 36, 37). Subsequently, NRF2 triggers antioxidant and antiapoptotic programs (20, 36), and promotes p62 upregulation, creating a loop that counteracts stress in autophagy-deficient conditions (36). We therefore infected BMDMs with *L. interrogans* serovar Manilae strain L495 to analyze NRF2 by immunofluorescence 4h post-infection. Whilst NRF2 staining was barely visible in the non-infected conditions, NRF2 staining in cell nuclei was clearly evident upon infection (**Fig 3A**). The intensity of NRF2 staining in the nucleus peaked around 4h post-infection (**Fig 3B**). We next performed single cell analysis of NRF2 fluorescence intensity in the nucleus of RAW264.7 cells infected for 4h with three different serovars of *L. interrogans*, and observed similar NRF2 increase for all the strains (**Fig 3C***, left panel*). Next, we calculated the ratio of NRF2 intensities [nucleus/cytoplasm] and estimated that cells positive for NRF2 had a ratio > 1.4. We then plotted the percentage of cells with NRF2 ratio superior to 1.4 and observed that up to 60% of cells were positive after 4h of infection (**Fig 3C***, right panel*). Finally, to understand the contribution of leptospiral LPS in triggering NRF2, we performed similar analyses on BMDMs stimulated for 4h with either live, heat-killed leptospires or their purified LPS. All conditions triggered similar NRF2 translocation (**Fig 3D**), suggesting that the leptospiral LPS, that triggers accumulation of p62, is responsible for NRF2 activation.

**Figure 3.**
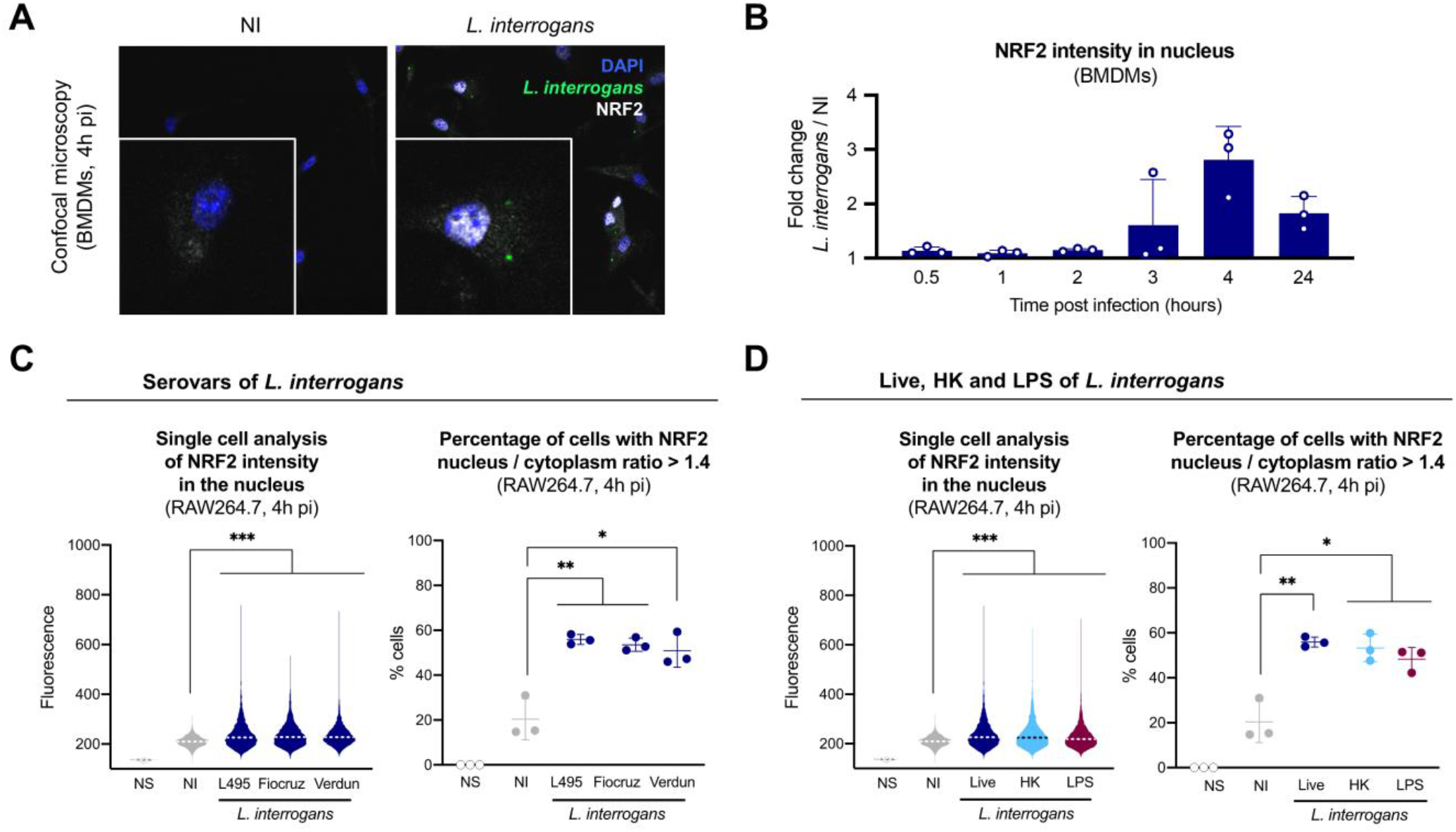
Leptospires and their LPS trigger translocation of transcription factor NRF2. **A)** Confocal microscopy analyses of NRF2 in BMDMs 4h post-infection with *L. interrogans* serovar Manilae strain L495 at MOI 100. **B)** Automated confocal microscopy analyses of NRF2 intensity 0.5h, 1h, 2h, 4h and 24h post-infection of BMDMs with *L. interrogans* serovar Manilae strain L495 at MOI 100. Fluorescence values of infected cells were normalized by non-infected cells. Bars correspond to mean +/- SD of technical replicates (*n*=3). **C) Left panel.** Single cell microscopy analysis of NRF2 intensity in the nucleus 4h post-infection of RAW264.7 cells with *L. interrogans* serovar Manilae strain L495, serovar Copenhageni strain Fiocruz L1-130 or serovar Icterohaemorrhagiae strain Verdun at MOI 100. Fluorescence is expressed in arbitrary units (AU). Dashed lines correspond to median. **Right panel.** Automated confocal microscopy analysis of the number of cells with NRF2 intensity ratio [nucleus/cytoplasm] > 1.4 4h post-infection of RAW264.7 cells with *L. interrogans* serovar Manilae strain L495, serovar Copenhageni strain Fiocruz L1-130 or serovar Icterohaemorrhagiae strain Verdun at MOI 100. Lines correspond to mean +/- SD of technical replicates (*n*=3). **D) Left panel.** Single cell microscopy analysis of NRF2 intensity in the nucleus 4h post-infection of RAW264.7 cells with MOI 100 of either live or heat-killed (56°C, 30 min) *L. interrogans* serovar Manilae strain L495 or stimulation with 1 µg/mL of the corresponding purified leptospiral LPS. Fluorescence is expressed in arbitrary units (AU). Dashed lines correspond to median. **Right panel.** Automated confocal microscopy analysis of the number of cells with NRF2 intensity ratio [nucleus/cytoplasm] > 1.4 4h post-infection of RAW264.7 cells with MOI 100 of either live or heat-killed (56°C, 30 min) *L. interrogans* serovar Manilae strain L495 or stimulation with 1 µg/mL of the corresponding purified leptospiral LPS. Lines correspond to mean +/- SD of technical replicates (*n*=3). **A-D)** Data presented are representative of at least 2 independent experiments.

### p62 and NRF2 are in a feedback loop upon infection with leptospires

Consequent to p62 accumulation and NRF2 translocation upon infection with *L. interrogans*, we investigated the potential regulation mechanisms between these two phenotypes. We performed small-interfering RNA (siRNA) transfection in BMDMs to knock-down specifically p62 or NRF2, and analyzed NRF2 and p62 4h post-infection with leptospires by automated microscopy. We observed that the NRF2 ratio [nucleus/cytoplasm] was lower upon infection in the p62 siRNA condition (**Fig 4A***, left panel*). Conversely, p62 puncta formation was reduced upon infection in NRF2 siRNA transfected BMDMs (**Fig 4B***, left panel*). Both p62 and NRF2 knocked-down were confirmed and quantified by automated microscopy (**Fig 4A** **& 4B,** *right panels*). Overall, these results showed that p62 contributed to NRF2 translocation, and that, in turn, NRF2 regulated p62 puncta formation, suggesting that p62 and NRF2 are in a feedback loop upon infection by *L. interrogans*.

**Figure 4.**
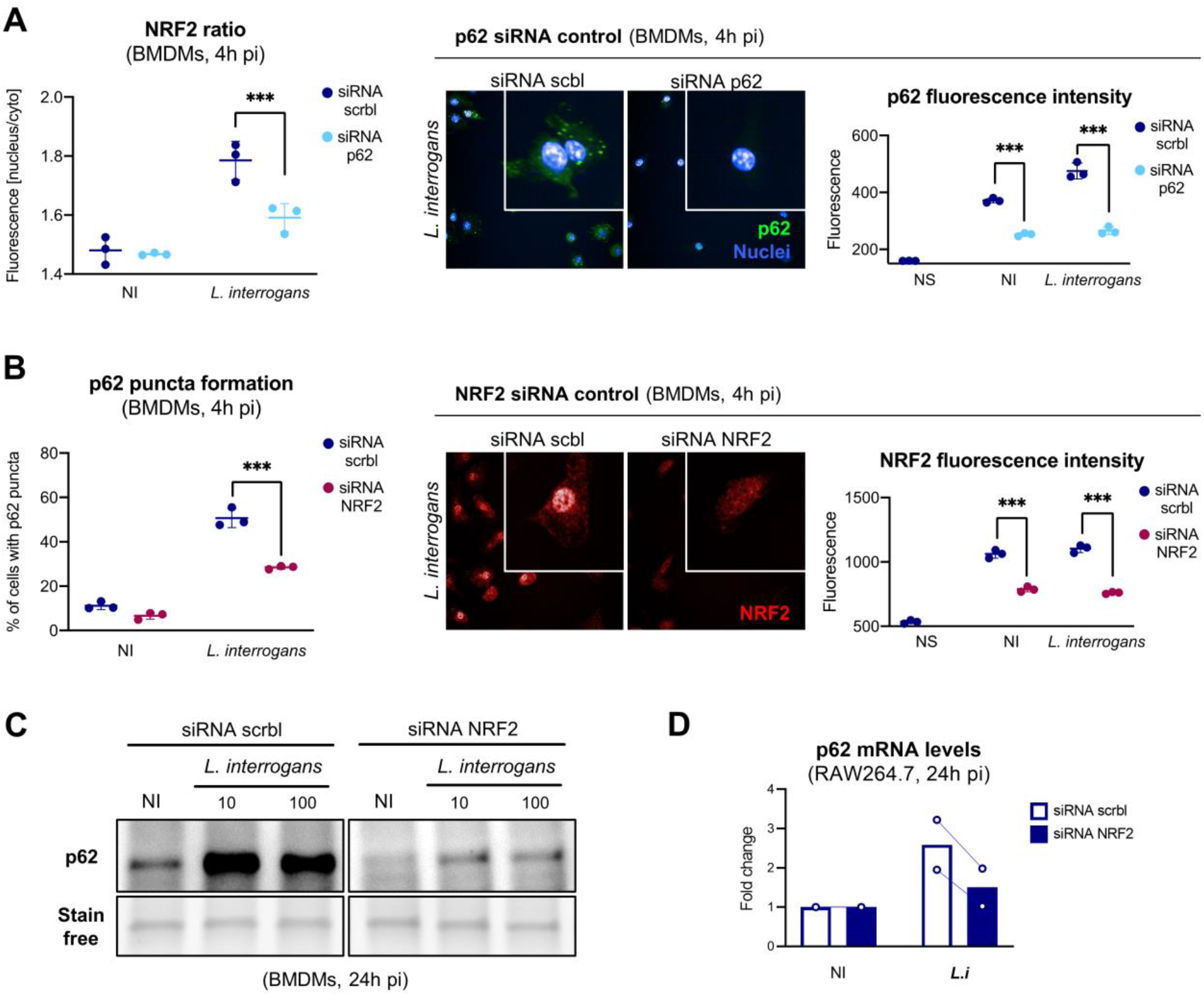
p62 and NRF2 are in a feedback loop upon infection with leptospires. **A) Left panel.** Automated confocal microscopy analysis of NRF2 intensity ratio [nucleus/cytoplasm] in BMDMs transfected with scramble (scrbl) or p62 siRNA and infected for 4h with *L. interrogans* serovar Manilae strain L495 at MOI 100. Lines correspond to mean +/- SD of technical replicates (*n*=3). **Right panel.** Corresponding controls are microscopy analyses and quantification of p62 intensity in the same transfection and infection conditions. Lines correspond to mean +/- SD of technical replicates (*n*=3). **B) Left panel.** Automated confocal microscopy analysis of p62 puncta formation in BMDMs transfected with scrbl or NRF2 siRNA and infected for 4h with *L. interrogans* serovar Manilae strain L495 at MOI 100. Lines correspond to mean +/- SD of technical replicates (*n*=3). **Right panel.** Corresponding controls are microscopy analyses and quantification of NRF2 intensity in the same transfection and infection conditions. Lines correspond to mean +/- SD of technical replicates (*n*=3). **C)** WB of p62 in BMDMs transfected with scrbl or NRF2 siRNA and infected for 24h with *L. interrogans* serovar Manilae strain L495 at MOI 10 or 100. **D)** RT-qPCR analyses of p62 mRNA levels in RAW264.7 cells transfected with scrbl or NRF2 siRNA and infected for 24h with *L. interrogans* serovar Manilae strain L495 at MOI 100. Bars correspond to mean of independent experiments (*n*=2). **A-B)** Data presented are representative of at least 3 independent experiments. **C-D)** Data presented are representative of 2 independent experiments. Statistical analyses were performed using Student’s *t*-test with corresponding *p* values: *** for *p* < 0.001.

To further characterize the involvement of NRF2 in p62 activation, we analyzed BMDMs transfected with NRF2 siRNA and infected with *L. interrogans* by Western blot and RT-qPCR 24h post-infection, allowing us to measure both the protein and mRNA levels of p62. Interestingly, we observed that NRF2 silencing prevented p62 protein accumulation (**Fig 4C**) but also reduced mRNA upregulation (**Fig 4D**), suggesting that NRF2 is a transcriptional regulator of p62 activation.

### NRF2 translocation prevents inflammation upon infection

NRF2 is a stress response transcription factor that promotes antioxidant and antiapoptotic programs upon activation (20, 36). Therefore, we monitored the transcription of targets genes involved in fighting oxidative stress in macrophages in response to infection. We infected BMDMs with *L. interrogans* and analyzed the levels of glutathione S-reductase, catalase and heme oxygenase 1 mRNA by RT-qPCR at 4 h and 24 h post-infection. Interestingly, we observed no upregulation of these transcripts (**Sup. Fig 4A, 4B & 4C**), suggesting that no antioxidant program is activated upon infection with *L. interrogans*.

NRF2 has also been shown to have repressor functions and to dampen inflammation by inhibiting the transcription of cytokines (38, 39), hence conferring resistance to inflammatory disease such as sepsis (38). We therefore investigated the role of NRF2 translocation in inflammation upon infection by leptospires. We transfected RAW264.7 cells with siRNA targeted against NRF2 and then infected them with *L. interrogans*. As expected, considering that NRF2 is known to prevent cell death, we observed a decrease in macrophage viability only upon infection of NRF2-silenced cells (**Fig 5A***, left panel*). Interestingly, this enhanced loss of viability was not associated with lactate dehydrogenase (LDH) release from the cytosol, illustrating that no membrane damage or cell lysis occurred (**Fig 5A***, right panel*). We then analyzed the mRNA levels of several cytokines and enzymes, namely iNOS (nitric oxide inducible synthase), IL6, TNF and IL10. Upon infection, all mRNA levels were higher in NRF2-silenced cells than in control cells (**Fig 5B**), suggesting that NRF2 does play a role in repressing the expression of inflammatory targets. Finally, to address whether this transcriptional regulation was strong enough to alter cytokines levels, we analyzed nitric oxide (NO), IL6, TNF and IL10 production in cell supernatant. Consistent with the mRNA analyses, we observed higher NO and TNF upon infection of NRF2-silenced RAW264.7 (**Fig 5C**). These findings were even more striking after normalizing cytokine production against cell viability, as measured using MTT (**Fig 5C***, grey areas*). Increased levels of IL6 were observed after normalizing for cell viability, whereas IL10 was not increased under any condition (**Fig 5C***, grey areas*). Overall, our results show that NRF2 plays a repressor role to dampen production of inflammatory mediators such as NO, TNF and IL6 in mouse macrophages upon infection with *L. interrogans*.

**Figure 5.**
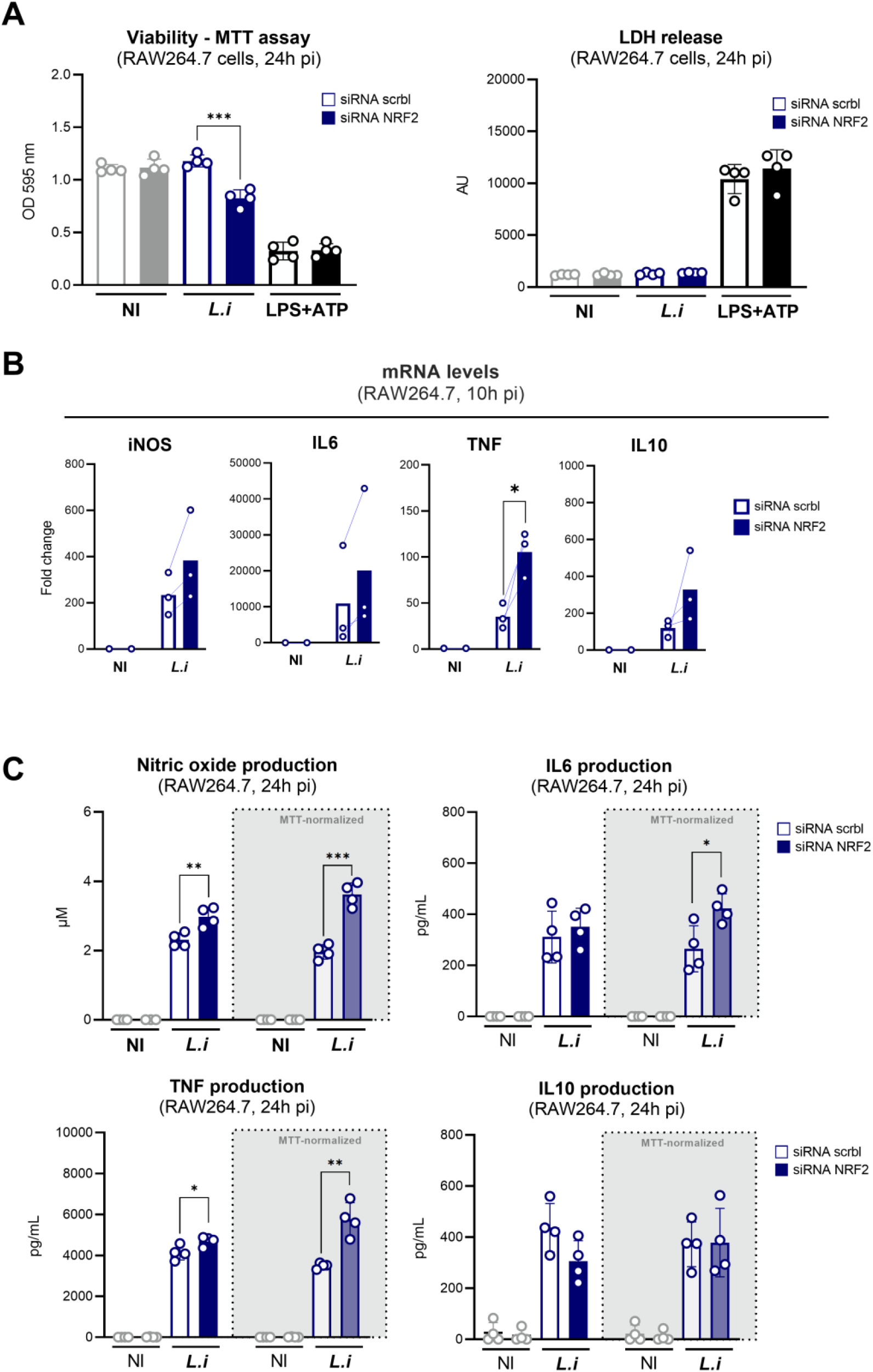
NRF2 translocation prevents inflammation upon infection. **A)** Cell viability (measured by MTT assay) and LDH release (measured by CyQuant assay) in RAW264.7 cells transfected with scramble (scrbl) or NRF2 siRNA and infected for 24h with *L. interrogans* serovar Manilae strain L495 at MOI 100 or stimulated with positive control for lytic cell death: LPS of *E. coli* 1µg/mL + ATP 5mM. Bars correspond to mean +/- SD of technical replicates (*n*=4). **B)** RT-qPCR analyses of iNOS, IL6, TNF and IL10 mRNA levels in RAW264.7 cells transfected with scrbl or NRF2 siRNA and infected for 10h with *L. interrogans* serovar Manilae strain L495 at MOI 100. Bars correspond to mean of independent experiments (*n*=3). **C)** NO dosage by Griess reaction and, IL6, TNF and IL10 dosages by ELISA in RAW264.7 cells transfected with scrbl or NRF2 siRNA and infected for 24h with *L. interrogans* serovar Manilae strain L495 at MOI 100. Bars correspond to mean +/- SD of technical replicates (*n*=4). **A-C)** Data presented are representative of at least 3 independent experiments. Statistical analyses were performed using Student’s *t*-test with corresponding *p* values: * for *p* < 0.05; ** for *p* < 0.01 and *** for *p* < 0.001.

## DISCUSSION

Our results evidenced an accumulation of the autophagy adaptor p62 in puncta-like structures in response to infection with both pathogenic *L. interrogans* and saprophytic *L. biflexa*. Classically, autophagy adaptor p62 accumulates on the surface of intracellular bacteria, such as *Salmonella* (*16*), *Listeria* or *Shigella* (40) to target them to autophagosomes and it is associated with active modulation of the autophagy flux through effectors (15, 16, 23, 27, 28). However, in the case of *Leptospira*, we have previously published that leptospires actively enter but do not remain inside macrophages and are not targeted by xenophagy (14). Moreover, the accumulation of p62 is also visible in response to stimulation with extracellular heat-killed leptospires or with purified LPS through TLR2/TLR4, hence excluding that only intracellular leptospires actively promote the formation of p62 puncta. This hypothesis is consistent with the fact that leptospires do not possess traditional secretion systems (41) used by other bacterial pathogens to modulate autophagy. Furthermore, our results showed that leptospires do not alter autophagy flux in macrophages. These findings are in accordance with previous studies from our laboratory showing that leptospires escape different innate immune autophagy-activating pathways (15, 42), such as NOD-like receptors (8) and TLR4-TRIF (13). Overall, we conclude that the leptospires behave differently from other pathogenic bacteria that actively modulate autophagy, and for which p62 accumulates on the surface of intracellular bacteria.

We further demonstrated that p62 accumulation was induced by leptospiral LPS signaling through TLR2/TLR4. The unique leptospiral LPS, the best characterized virulence factor (10), plays an important role in leptospiral host-pathogen interactions. We previously showed that leptospiral LPS avoids human TLR4 and mouse TLR4-TRIF activation (13). Recently, leptospiral LPS has also been shown to prevent cell death by pyroptosis (43). Therefore, it was not surprising to find the leptospiral LPS responsible for the p62-puncta phenotype. Interestingly, the LPS of *L. biflexa* has been shown to be a more potent TLR4 agonist than the LPS of pathogenic *L. interrogans* (13), consistent with our results showing greater p62 accumulation observed after infection with the saprophytic strain. Overall, these findings are consistent with other studies reporting that p62 accumulates *via* TLR signaling (44), directly through NFκB-mediated transcriptional regulation. We believe this could be the case upon infection with leptospires because: *(i)* our results show an upregulation of p62 mRNA upon infection and *(ii)* p62 levels upon infection are much higher than upon autophagy blockage with BafA1, suggesting that the accumulation of p62 is not mediated by a dysregulation of autophagy.

TLR-induced accumulation of p62 has been shown to trigger the translocation of the transcription factor NRF2 (44). Under physiological conditions, NRF2 is classically targeted for degradation by the proteasome *via* interaction with its inhibitor Keap1 (37, 45). The accumulation and direct binding of p62 to Keap1, leads to its sequestration and subsequent release from NRF2, which then can translocate in the nucleus (20, 36, 37, 45). Our results confirmed that p62 induces translocation of NRF2 upon infection with leptospires, and illustrated that the feedback loop between p62 and NRF2, described in the literature (36), is activated upon *L. interrogans* infection. Interestingly, this loop has few negative regulators, and it is hypothesized that its main regulator would be autophagy, through degradation of p62 (36). Therefore, we hypothesize that since infection with leptospires does not trigger autophagy, the feedback loop between p62 and NRF2 remains active, explaining why p62 puncta did not disappear even at 24h post-infection. Overall, out data indicate that leptospires are potent activators of the p62/NRF2 axis in macrophages.

Interestingly, NRF2 has also been shown to have transcription inhibition properties. In BMDMs, NRF2 was described as an inhibitor of transcription for pro-inflammatory targets such as IL6, IL1β and IL12 (39, 46). Our results showed that this is also the case upon infection with *L. interrogans*. NRF2 inhibits polymerase III recruitment and hence prevents transcription of cytokines in response to stimulation (39). Consequently, NRF2 was shown to downregulate neutrophils activation and migration (47). Interestingly, although neutrophils are abundant in the blood of mice and humans infected with leptospires, they are barely observed in the kidneys, the niche occupied by leptospires during chronic infection (48, 49). Whether *Leptospira* modulate NRF2 translocation in neutrophils *via* their LPS to favor survival in the kidneys remains to be studied. In addition, we showed that the accumulation of p62 observed in murine macrophages was conserved in human cells. Interestingly, human cells do not sense leptospiral lipid A through TLR4 (12), leading us to hypothesize that TLR2 alone could be responsible for sensing leptospires and inducing p62 accumulation in THP1 cells. Although we could not address NRF2 translocation in human cells due to lack of specific tools, we speculate that inflammation dampening might be conserved in human macrophages.

Other microbes such as Epstein-Barr virus (EBV) or parasite *Leshmania major* have been shown to activate the p62/NRF2 axis (50, 51), hence showing an important role of NRF2 in response to pathogens. However, to date, the role of NRF2 translocation in response to pathogens remains unclear. Among others, NRF2 is involved in the induction of antioxidant program upon infection (36). We were therefore surprised to observe no modulation in the mRNA levels of antioxidant targets upon infection. Of note, pathogenic leptospires are equipped to fight against oxidative stress with inducible catalase, a virulent factor, peroxidase, and peroxiredoxin (52). Interestingly, active repression of these NRF2-dependent antioxidant targets was previously shown upon infection with live *Leshmania major* parasite (51, 53). Whether leptospires could also actively prevent upregulation of NRF2 antioxidant targets remains to be addressed.

In summary, we have demonstrated that leptospires passively subvert the p62-NRF2 axis through LPS activation of TLR signaling and this leads to a reduction of inflammatory mediators. This original mechanism might play a key role in the discretion of leptospires in hosts, evasion of immune cell recruitment, and could potentially contribute to the establishment of chronic infections.

## ACKNOWLEDGMENTS

We acknowledge Nathalie Aulner and Anne Danckaert (Institut Pasteur) as well as the Photonic BioImaging Unit of Technology and Service, a member of the France-BioImaging infrastructure, for support in conducting this study. We thank Richard Wheeler (Institut Pasteur) for critical reading of the manuscript and English editing.

## STATEMENT OF ETHICS

All protocols were undertaken in compliance with EU Directive 2010/63 EU and the French regulation on the protection of laboratory animals issued on February 1^st^, 2013. They are part of project number # 2014-0049 and HA0036, which were approved by the Institut Pasteur ethics committee for animal experimentation (Comité d’Ethique en Expérimentation Animale CETEA registered under #89) and was authorized under #8562 by the French Ministry of Research, the French Competent Authority.

## AUTHORS CONTRIBUTIONS

Conception, administration, and supervision of the project: CW. Investigation: DB, IS, SP and CW.

Methodology: IS, DB and PE. Data analysis: DB, IS, PE and CW. Visualization: DB.

Validation: IS, DB, SP and CW. Resources: FVP.

Funding acquisition: CW and IGB.

DB and IS wrote the original draft, under CW’s supervision.

All authors contributed to the review of the manuscript and approved it for submission.

## DATA AVAILABILITY STATEMENT

All data generated or analysed during this study are included in this article. Further enquiries can be directed to the corresponding author.

---

**Sup. Figure 1.**
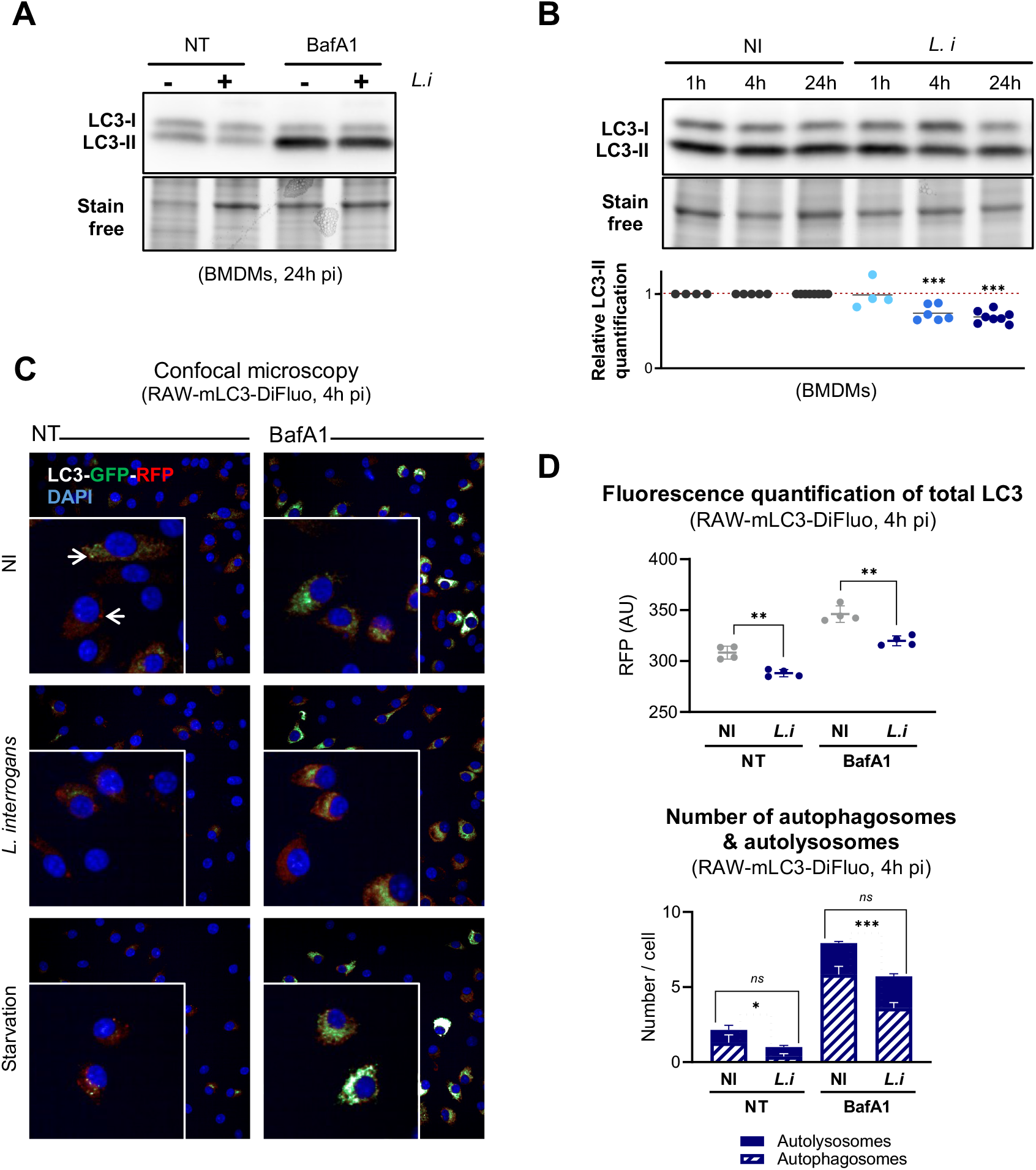
*L. interrogans* does not induce autophagy in murine macrophages and reduces the number of autophagosomes. **A)** WB of LC3 in BMDMs 24h post-infection with *L. interrogans* serovar Manilae strain L495 at MOI 100 with or without addition of bafilomycin A1 (BafA1) 4h prior to cell collection. **B)** WB and quantification of LC3 in BMDMs 1h, 4h or 24h post-infection with *L. interrogans* serovar Manilae strain L495 at MOI 100. Quantifications represent the ratio of LC3-II in stimulated cells over non-stimulated cells, and after gel stain free normalization. Each dot corresponds to the quantification of a WB from an independent experiment. **C-D)** Automated confocal microscopy images and quantifications of RAW-mLC3-DiFluo cells 4h post-infection with *L. interrogans* serovar Manilae strain L495 at MOI 100, or post starvation induction in EBSS medium, with or without BafA1 addition 2h before cell fixation. Of note, RAW-mLC3-DiFluo cells are stably transfected with LC3 tagged with tandem GFP-RFP. Both GFP and RFP are fluorescent under basal conditions, however, GFP fluorescence is quenched during acidification of autophagosomes upon fusion with lysosomes. Total LC3 fluorescence was measured based on RFP signal. Autophagosomes were quantified as RFP^+^/GFP^+^ punctas (upper white arrow) whereas autolysosomes were quantified as GFP^-^/RFP^+^ punctas (lower white arrow). Lines and bars correspond to mean +/- SD of technical replicates (*n*=4). **A-D)** Statistical analyses were performed using Student’s *t*-test with corresponding *p* values: * for *p* < 0.05; ** for *p* < 0.01 and *** for *p* < 0.001.

**Sup. Figure 2.**
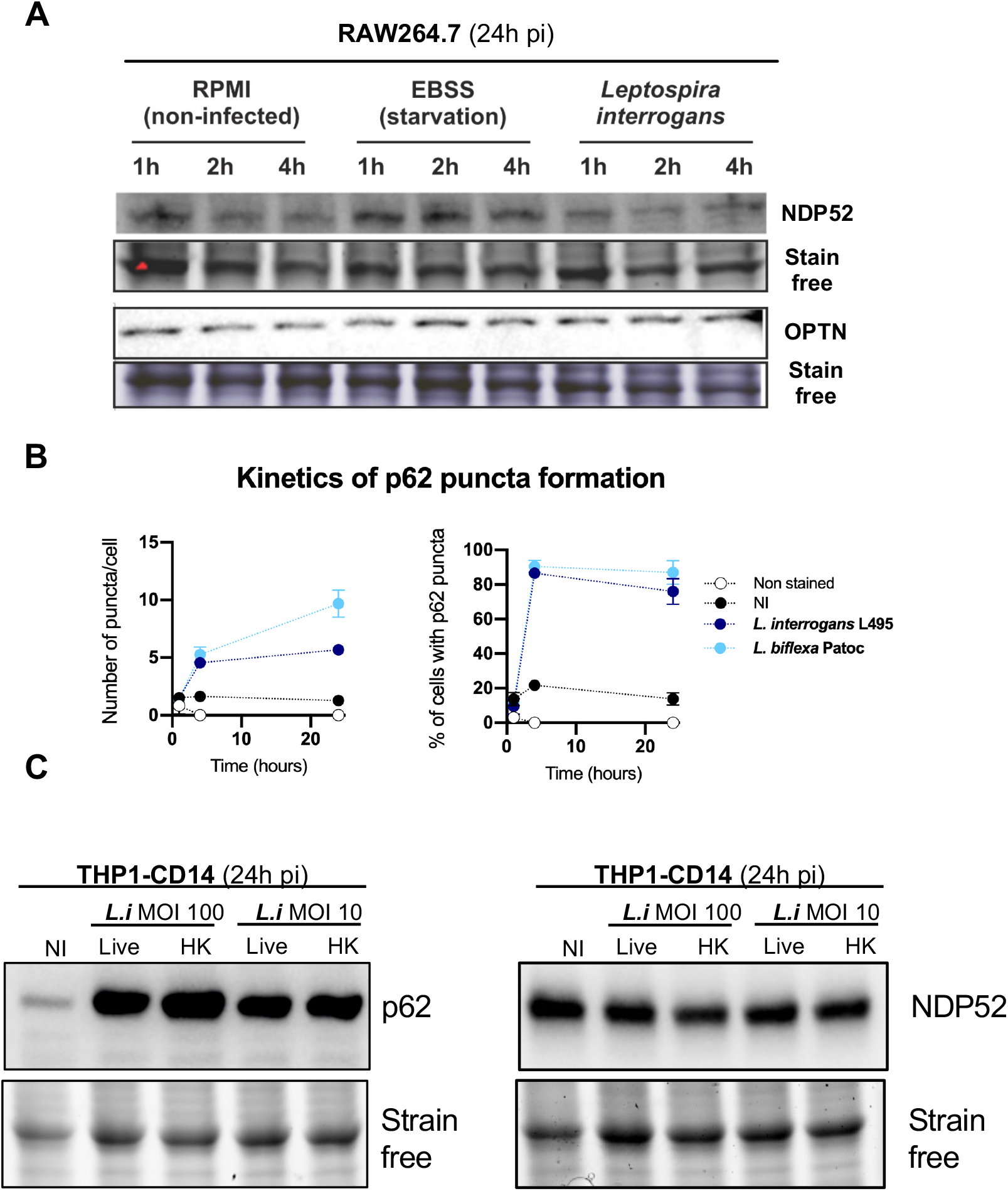
Specific p62 accumulation is observed in murine and human cells. **A)** WB of NDP52 and OPTN in RAW264.7 cells 1h, 2h and 4h post-infection with *L. interrogans* serovar Manilae strain L495 at MOI 100, or upon starvation induction in EBSS medium. **B)** Automated microscopy analyses of the number of p62 puncta *per* cell and the percentage of p62 positive cells 1h, 4h and 24h post-infection of RAW264.7 cells with *L. interrogans* serovar Manilae strain L495 or *L. biflexa* serovar Patoc strain Patoc I, at MOI 100. Dots correspond to mean +/- SD of technical replicates (*n*=3). Statistical analyses were performed using Student’s *t*-test with corresponding *p* values: * for *p* < 0.05; ** for *p* < 0.01 and *** for *p* < 0.001. **C)** WB of p62 and NDP52 in human THP1-CD14 cells 24h post-infection with *L. interrogans* serovar Manilae strain L495 at MOI 100.

**Sup. Figure 3.**
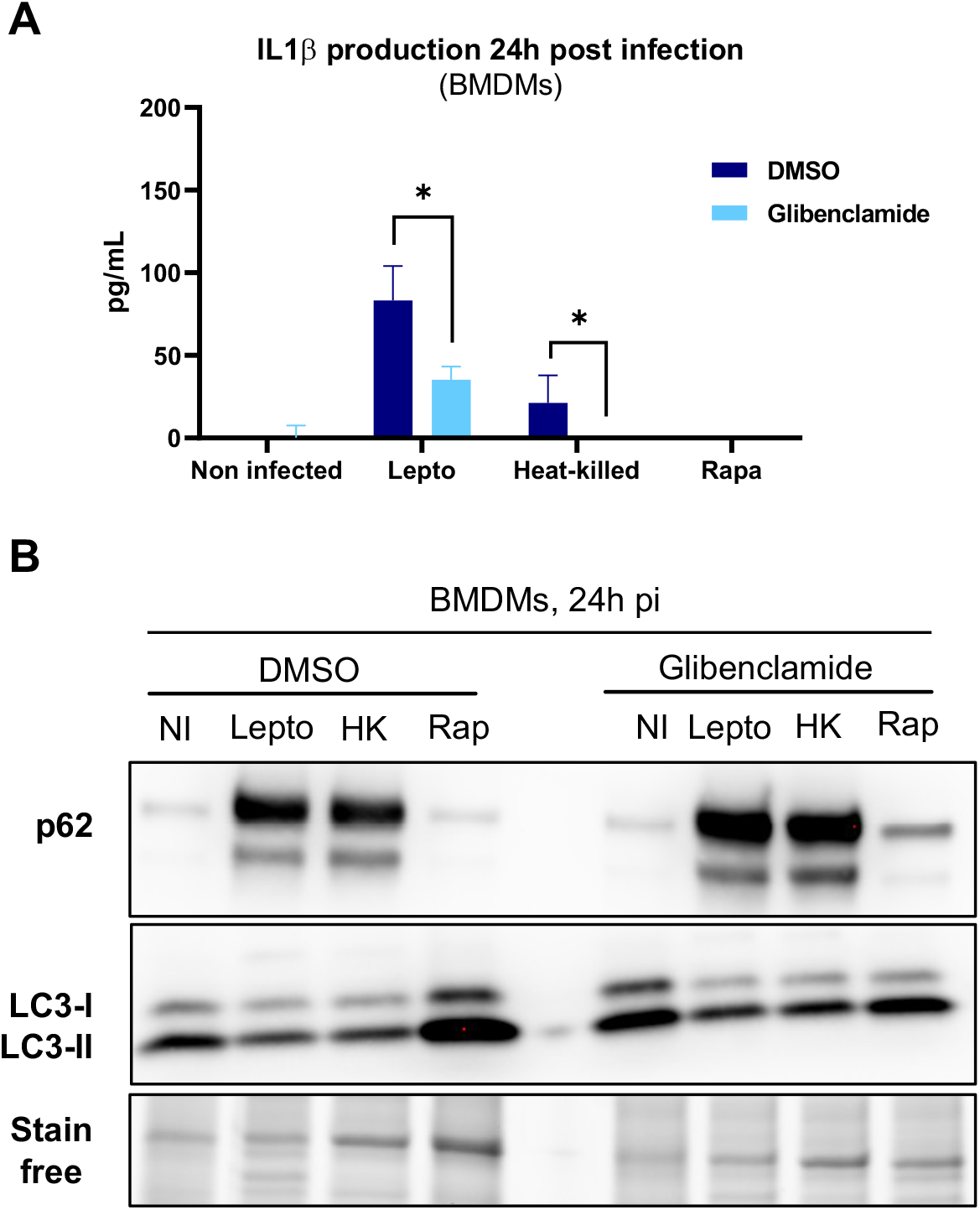
p62 accumulation induced by leptospires does not rely on NLRP3 inflammasome. **A)** IL1β dosage by ELISA in BMDMs 24h post-infection with either live or heat-killed (56°C, 30 min) *L. interrogans* serovar Manilae strain L495 at MOI 100 with or without 25 µM glibenclamide (inflammasome inhibitor). Bars correspond to mean +/- SD of technical replicates (*n*=4). Statistical analyses were performed using Student’s *t*-test with corresponding *p* values: * for *p* < 0.05. **B)** WB of LC3 and p62 in BMDMs 24h post-infection with either live or heat-killed (56°C, 30 min) *L. interrogans* serovar Manilae strain L495 at MOI 100, or upon autophagy induction with 500 mM rapamycin (Rap) with or without 25 µM glibenclamide.

**Sup. Figure 4.**
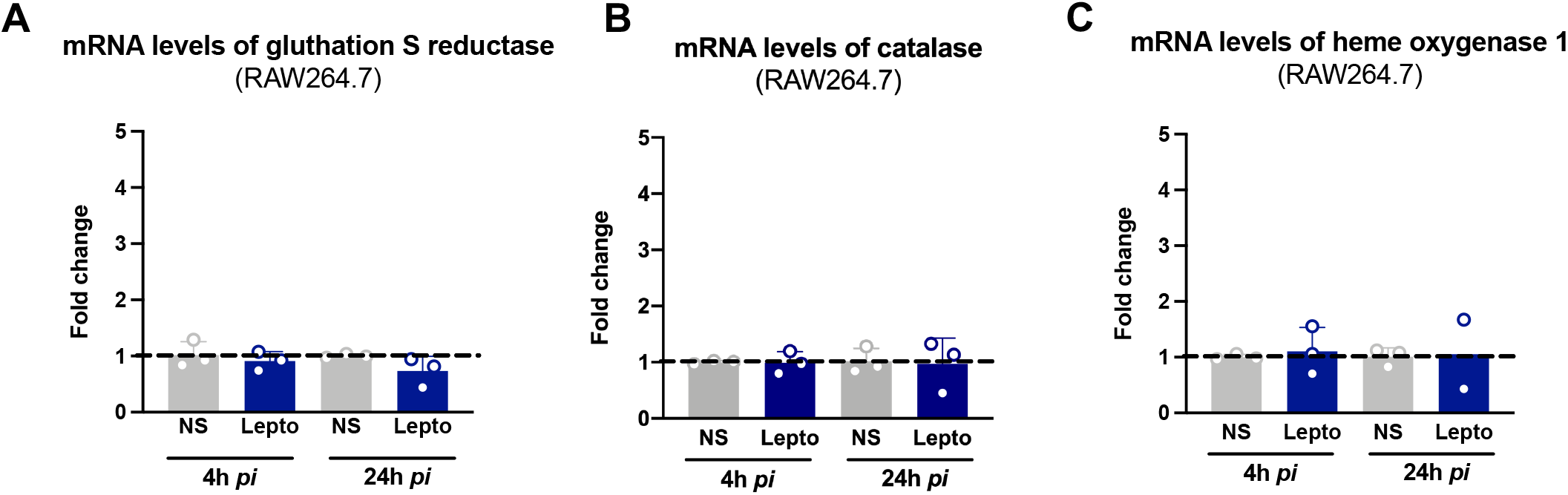
Infection by leptospires does not trigger transcription of NRF2 antioxidant targets. **A-B)** RT-qPCR analyses of gluthatione S-reductase, catalase and heme oxygenase 1 mRNA levels in RAW264.7 cells 4h or 24h post-infection with *L. interrogans* serovar Manilae strain L495 at MOI 100. Bars correspond to mean +/- SD of technical replicates (*n*=3).

